# Singular superlet transform achieves markedly improved time-frequency super-resolution for separating complex neural signals

**DOI:** 10.1101/2023.02.27.530211

**Authors:** Kaan Kesgin, Henrik Jörntell

## Abstract

Time-frequency decomposition is a well-established method to unmix signals generated by multiple sources with unique characteristics. However, there are cases of high signal complexity where existing time-frequency decomposition tools are insufficient for localizing and representing short-bursting signals. One example is the currently highly popular extracellular low-impedance recordings from multi-electrode arrays in the brain *in vivo* where each neuron repeatedly generates a specific signal ‘fingerprint’ (characteristic spike waveform) that can be mixed with the signals of 100s of other sources, including the spikes of nearby neurons. Here we derive the singular superlet transform (SST) method, which enables highly localized representations of fast and short bursts compared to other super-resolution spectral estimators, while also requiring orders of magnitude fewer operations. We demonstrate a substantial edge of SST over current methods in isolating specific neuronal spikes with high-fidelity in challenging, complex recording signals from neocortex *in vivo*. We also exemplify SST’s generic signal processing capability by achieving outstanding resolution in the decomposition of complex acoustic data.

## 1 Introduction

In neuroscience, the aim to record from larger and larger populations of separable neurons is driving a dramatic hardware development of multi-electrode arrays and brain-machine interfaces. For such systems, lower electrical impedance is a key design feature [1–6] as it widens the ‘field of view’ of each electrode. This comes with several advantages such as allowing for a larger yield of captured neurons, resistance to movement artefacts and even relative reduction of labor for performing the experiments as the electrode no longer needs to be placed in the immediate vicinity of a neuron [7]. This increased field of view per electrode, however, creates a need for dense recording arrays and more sophisticated algorithms to achieve accurate spike clustering [8–11]. The algorithmic challenge is analogous to the cocktail party problem: speech sounds from different individuals mix in the air before arriving at the ear. In order to follow at least one conversation, we then need to unmix the sound signatures to estimate their individual sources. This is possible if those sources have unique time-frequency ‘fingerprints’. Here we propose such time-frequency decomposition to be highly advantageous also for the unmixing of neural recording signals.

A standard approach to calculate the time-frequency spectrum of a signal is the short-time Fourier transform (STFT) [12], which uses a fixed sliding window to compute the Fourier transform of the time-continuous signal. An issue with STFT is that if the window size is chosen to capture slower components of extracellular events (~ 1-500 Hz) [13] with high enough resolution, it would be too broad to capture faster bursts (~ 1000-8000 Hz) with good resolution. This limits STFT from being applicable to unmix extracellularly recorded neuronal spikes, which typically span a wide frequency range [14–16]. A multi-resolution method that can better address this issue is the continuous wavelet transform (CWT) [17], based on the good time-frequency tuning characteristics of the Morlet wavelet [18]. CWT achieves good temporal resolution across scales by uniformly compressing the wavelet (ie: the window) as a function of the frequency of interest. However, due to the Heisenberg uncertainty principle, there is a fundamental trade-off between localizing the signal accurately in the time domain versus the frequency domain [19]. Hence, in the case of CWT, as the wavelet is compressed in time with respect to increased frequency, the frequency resolution becomes poor.

In order to limit the trade-off effects in such singular observation spectrograms, combining multiple spectrograms with different window sizes has been proposed [20, 21]. Methods where multiple observations are combined, typically using a geometric mean operation, has been named “super-resolution” methods [22, 23]. The superlet transform (SLT) is a recent multiple observations method, which is derived from CWT, but combines observations of multiple wavelets into one. SLT has been shown to perform well in localizing low frequency bursts in electroencelographic (EEG) data [24]. However, as we will show, for the higher frequencies in a wide frequency dynamic range, SLT experiences a similar problem as CWT: the high frequency resolution gets smeared when combined with high temporal resolution observations. But since SLT also has a drastically increased computational load compared to CWT, this effect alone renders SLT impractical for investigations of large data sets [25] especially when the signals involve a wide dynamic frequency range, such as large scale extracellular neural recording data [26].

Here, we build on the principles of the SLT [24], but rather than relying on calculating multiple discrete observations across resolutions to generate their mean, we instead derive a singular mathematical expression of a wavelet, called the singular superlet, to generate a similar fundamental effect. We show that using this superlet inside a wavelet transform, the singular superlet transform (SST), achieves super-resolution time-frequency decomposition while also mitigating both the wide-dynamic range resolution issue and the issue of the computational load in existing super-resolution methods. We investigate the applicability of SST to identify neuronal spike signal ‘fingerprints’ in intra-cortical low-impedance multi-electrode array recordings *in vivo* (Fig.1a,b). We compare SST’s resolution to those of CWT and SLT (Fig.1c), which are both based on Morlet wavelets (Fig.1d) and are closely related to our singular superlet (Fig.1,e).

**Fig. 1.**
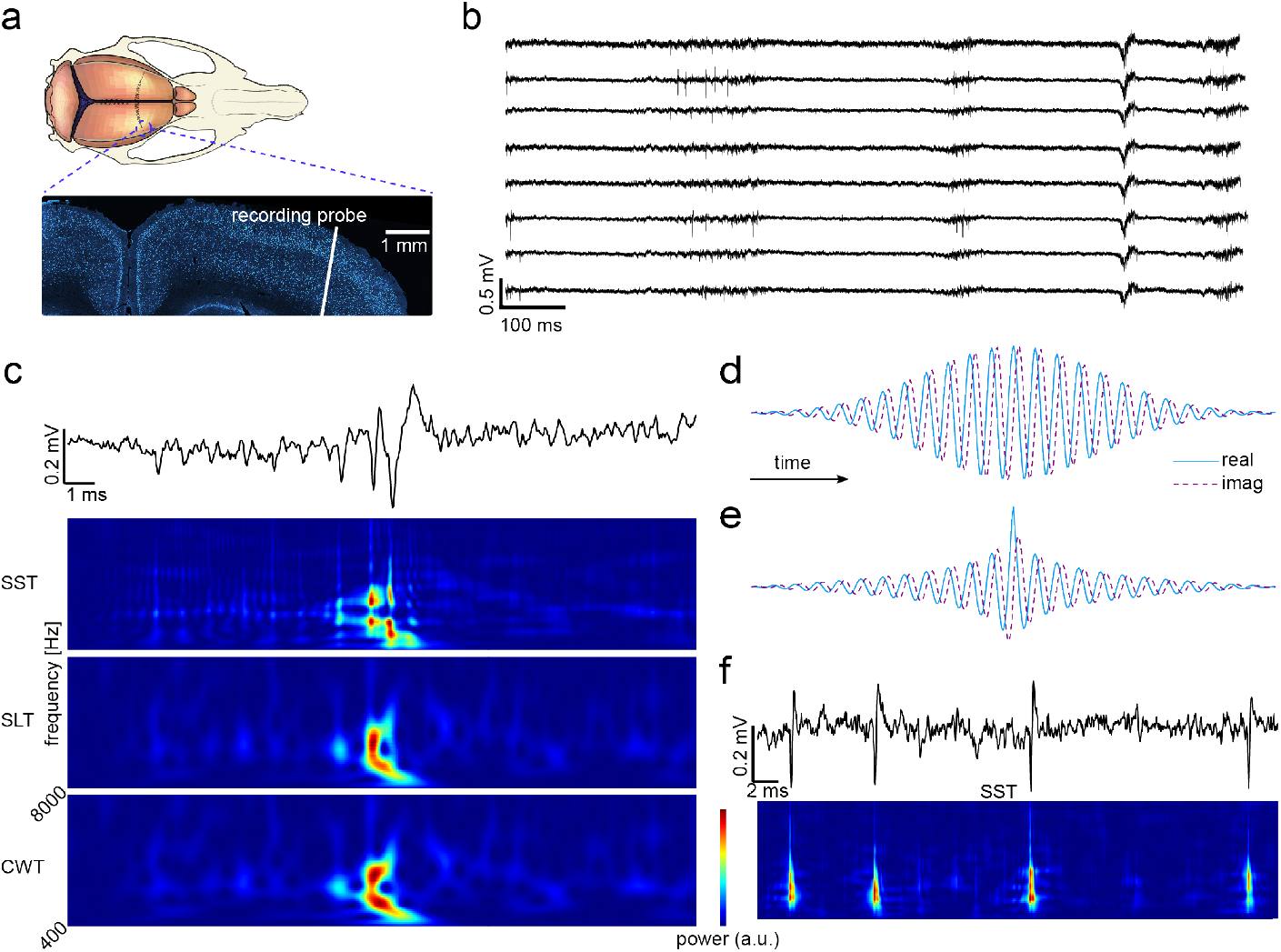
Spike recording data and remapping to the time-frequency domain. **a.** Recording location in a transverse histological section of the cortex. White line represents the location of the NeuroPixels probe multielectrode array. **b.** Example of time-continuous data across 8 neighboring electrodes during an intracortical recording. **c.** Zoomed in time-continuous data showing spiking events from the cortex and how these events appear in SST as well as in the two comparative time-frequency decomposition methods SLT and CWT. **d.** Complex Morlet wavelet used commonly in CWT and SLT with its temporal representation. **e.** Representation of the singular superlet (used in the present SST method). **f.** Time-continuous trace of spikes repeatedly generated by the same neuron (i.e. one cluster of spikes), and its singular superlet transform (SST). Note the high time-frequency resolution and the reproducibility of the signal ‘fingerprint’ across the 4 sequential spikes in SST.

We further compare SST to band-pass filtering (BPF), which is widely used in current state-of-the-art methods for spike sorting [2, 3, 10, 27–35]. BPF is utilized to facilitate spike detection, but as we will demonstrate BPF can potentially be problematic as the interferences between the multiple independent signal sources which are normally present in brain recordings can distort the information obtained after BPF, an effect that is amplified with low-impedance electrodes. This can in turn lead to spikes being missed or spuriously generated, or spike shapes becoming distorted and therefore misclassified. We find that by avoiding the distortion in the filtering step, SST provides both much more reliable spike detection and feature extraction, and therefore more reliable spike clustering. Apart from providing a promising basis for spike detection and neuronal clustering algorithms for multi-electrode array recording data, we also illustrate the multi-disciplinary use cases for SST applied to acoustic data, which results in a drastically improved time-frequency resolution over state-of-the-art, CWT-based, scalograms.

## 2 Results

### 2.1 The singular superlet

In order to understand the singular superlet transform (SST) we work our way up from a more intuitive super-resolution time-frequency decomposition method, the superlet transform (SLT) [24]. SLT works as follows; for a frequency of interest *f_c_*, multiple discrete observations on the input vector *x*(*t*) are combined geometrically or arithmetically as:

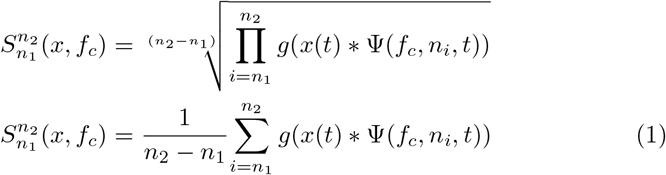

Ψ(*f_c_, n_i_, t*) is the modified complex Morlet wavelet [18] (figure 1-d) for central frequency *f_c_* and number of cycles *n_i_*, expressed as:

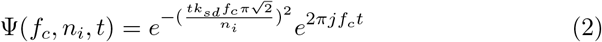

where *k_sd_* is the standard deviation of the Gaussian envelope and usually is kept constant (*k_sd_* = 5). The term *g*(*x*) = 2|*x*|^2^ in Equation 1 is due to convention and doesn’t impact the analysis on the super-resolution aspect of the underlying mean-of-convolutions operation. Hence it will be omitted in the derivation of the SST.

Going beyond SLT, we begin with considering the arithmetic mean operation for the sake of simplicity. In Equation 1, the discrete increments of *n_i_* can be taken infinitesimally small (*dn* → 0) to express the arithmetic mean operation from Equation 1 as:

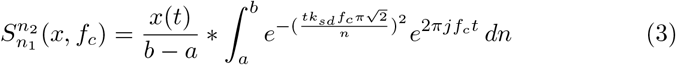

Focussing on the integral in Equation 3, in effect we create an integrated wavelet from Ψ(*f_c_, n, t*), consisting of infinitesimally small increments in the number of cycles of the wavelet. Though in principle the solution to this integral cannot be expressed in terms of fundamental operations, the definite integral has the form:

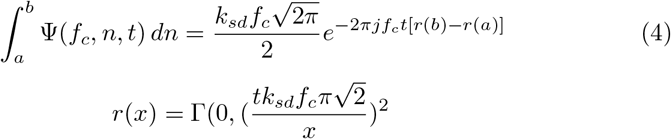

where Γ(0, *x*) is the lower incomplete gamma function.

In order to get an intuitive understanding of what captures the temporal behavior of this function in terms of fundamental operations we compute the series expansion for the indefinite integral from Equation 4 at n=0

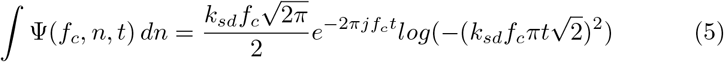

This shows that wrapping the oscillatory behavior in time around a *log*(−*t*^2^) envelope (or a *log*(*t*^2^) by symmetry) is the key to achieve time-frequency super-resolution using a singular wavelet transform per frequency of interest. Based on this observation we propose the singular superlet as:

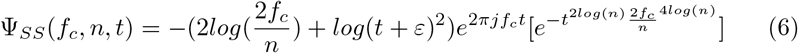

where 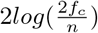 in Equation 6 is scaling the log decay in time to 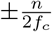 (in seconds, assuming central frequency *f_c_* is in Hz and *t* is in seconds). Intuitively, the smaller 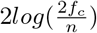 is, the higher the time resolution. Similarly, *n* is the number of cycles and is analogous to the case for the Morlet wavelet, the higher this number is, the better the frequency resolution. *ε* is a correction factor in the log envelope to ensure 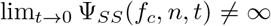 and empirically taken as half of the sampling interval. Finally the square bracketed terms are to ensure that the wavelet decays to zero in time towards infinity, however in practice one can also omit the square bracketed terms and cut the wavelet between 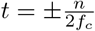, then zero pad to the desired length in time.

Following Ψ_*SS*_(*f_c_*, *n, t*), the singular superlet transform at the frequency of interest *f_c_* can be implemented as:

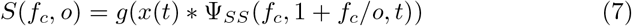

where *o* is the step parameter that adaptively adjusts the number of cycles in the singular superlet set to achieve uniform frequency representation as the frequency increases. Inside a transform, Ψ in above equations are normalized as described in Methods.

### 2.2 Applying SST to synthesized data

We explored the resolution of the SST time-frequency decomposition using synthesized signals (Fig.2). We compared the resolution only with CWT and SLT as these two methods have been applied to analyze neural data and they have also been extensively compared with other known time-frequency decomposition methods in [24] where they were found to have a performance edge over these other methods. Secondly, they are both wavelet based methods with identical parameters to SST and changing the parameter settings will hence have the same effect on all three methods. The first synthesized input vector (signal) was ‘overlaid chirps’ (Fig.2a), where a fast chirp from 250 Hz to 2750 Hz and a slow chirp from 850 Hz to 1800 Hz are linearly summed (see Methods). ‘Overlaid chirps’ was designed for a simple verification of the frequency behavior of the time-frequency decomposition. Although the three methods appeared to perform similarly, closer inspection reveals that the SST provided a higher resolution in several phases of the overlaid chirps. But the higher resolution of the SST became more evident for the ‘neighboring burst’ signal (Fig.2b). The initial burst consisted of two Morlet wavelets (see equation 2) with the central frequency *f_c_* = 300 Hz and number of cycles *n_i_* = 3, overlapping for 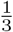 of their duration. The middle burst consisted of two Morlet wavelets where *f*_*c*_1__ = 1250 Hz and *f*_*c*_2__ = 1725 Hz with *n_i_* = 6 for both, overlapping in time for 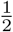 of the 1725 Hz wavelet’s duration. The final burst was a sine wave of 5 cycles at 2300 Hz. The middle burst singled out SST as being capable of separating the two wavelets, whereas they became conjoined both with CWT and SLT. The last synthetic signal was a ‘neural recording-like’ test scenario to explore how randomly scattered, overlaid high-frequency bursts, with random number of cycles, were resolved (Fig.2c). Notably, the peak of each cycle (dashed lines) could be distinguished only with SST.

**Fig. 2.**
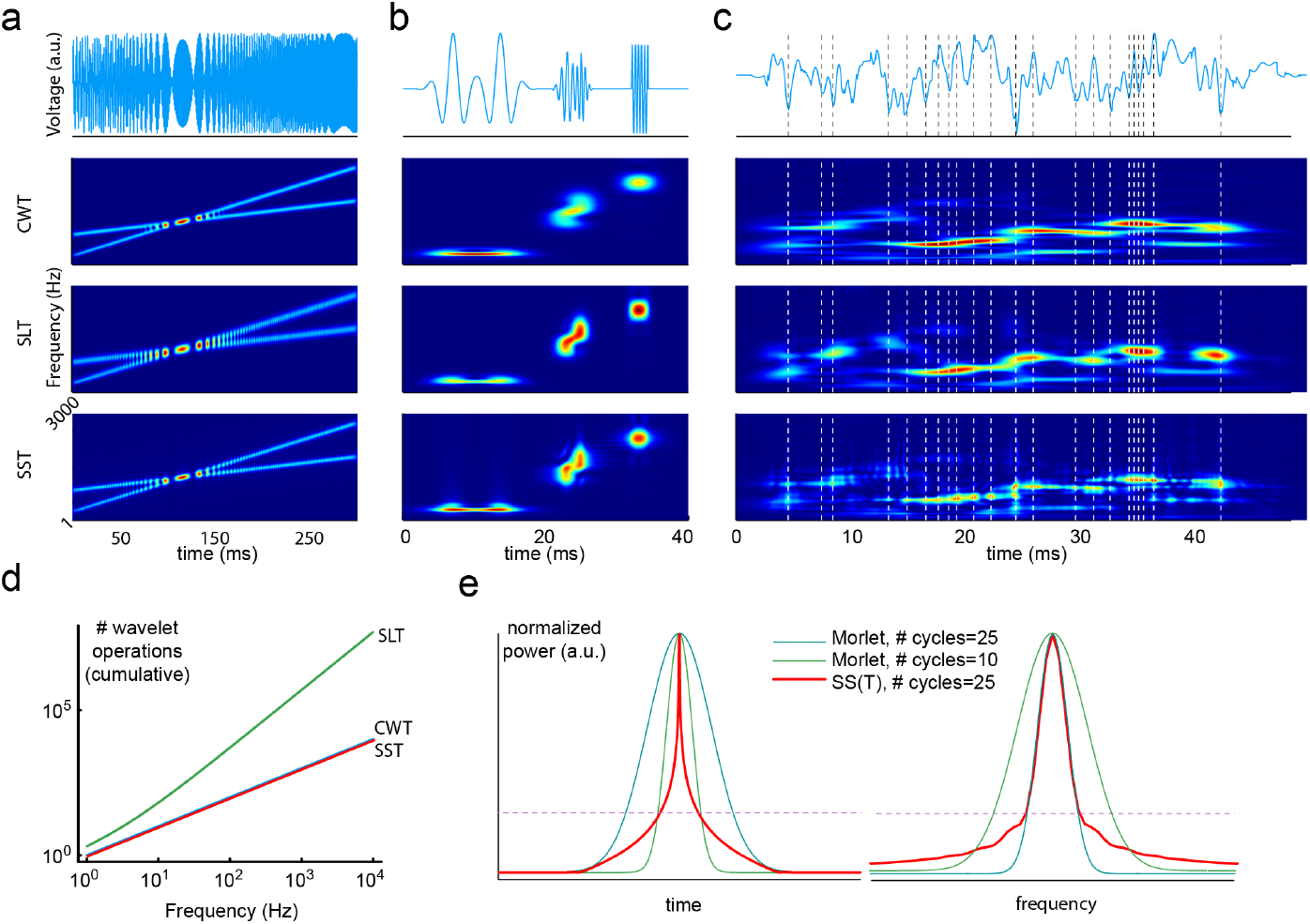
Performance comparison for synthesized signals with complex timefrequency characteristics. **a** Synthesized ‘overlaid chirps’ signal (top) and time-frequency decompositions achieved by CWT, SLT and SST, respectively. **b.** Similar display as in **a** for the synthesized ‘neighboring burst’ signal. **c.** Similar display as in **b** for the synthesized ‘random burst’ signal. **d** number of cumulative operations required for each given method with respect to frequency. **e** Resolution comparison for wavelets in time and frequency spaces at 3500 Hz (*f_c_*) for Morlet wavelets (used in CWT and SLT) with different number of cycles (*N_c_*) and for a Singular Superlet (used in SST). Dashed line is located at 25% of the peak power and all three wavelets are chosen to have the same value around that point.

The higher resolution of SST did not come with the trade-off of requiring a higher number of windowed operations to isolate the peaking events (Fig.2d), which is the case with other super-resolution methods such as SLT. As the derivation of the SST at the logical level involves an analytic expression of infinite observations with continuous, infinitesimally small increases in window sizes, that effect is technically captured within a single wavelet observation using SST, making the computational load of SST much below the super-resolution SLT and exactly on-par with the traditional CWT (Fig.2d).

The Heisenberg uncertainty principle dictates that an apparent increase in resolution in either the time or the frequency domain will come with a tradeoff. It is important to know where this trade-off occurs, as this may impact the choice of method for a specific problem at hand. Constructing the marginals for the singular superlet, and the Morlet wavelet for comparison, illustrates how the perceived time-frequency super-resolution with SST is achieved (Fig.2e). Around the central points of interest in time and in frequency, respectively, most of the represented power of the singular superlet ends up being sharper than a higher-temporal-resolution Morlet wavelet in the time domain and as sharp as a higher-frequency-resolution Morlet wavelet in the frequency domain. Hence, the trade-off for the singular superlet is the increased uncertainty at the tails away from the central points of interest, in both domains. SST is hence well-suited for resolving high-power bursts like extracellular spiking data, at the cost of potentially lower performance for uniformly confining long cycles within narrow frequency bands.

### 2.3 Applying SST to neuronal spike data

In low-impedance multi-electrode recordings from the neocortex (Fig.1a), the signal in each recording electrode represents the combined activity of all neurons, as well as the field potentials, that are within the field of view of the electrode (Fig.1b). In order to achieve an accurate spike sorting from this type of complex neural data, sorting algorithms break down the problem in multiple steps: data filtering, spike detection, spike feature extraction and clustering. Since they are structured sequentially, the quality of the information extracted from one step impacts the success of the subsequent steps, where the over-arching goal is to interpret the raw biological information with the highest fidelity. Therefore, we compared the levels of distortion of the raw biological data caused by band-pass filtering (BPF) and SST to investigate the applicability of SST in the filtering stage of spike sorting (figure 3).

**Fig. 3.**
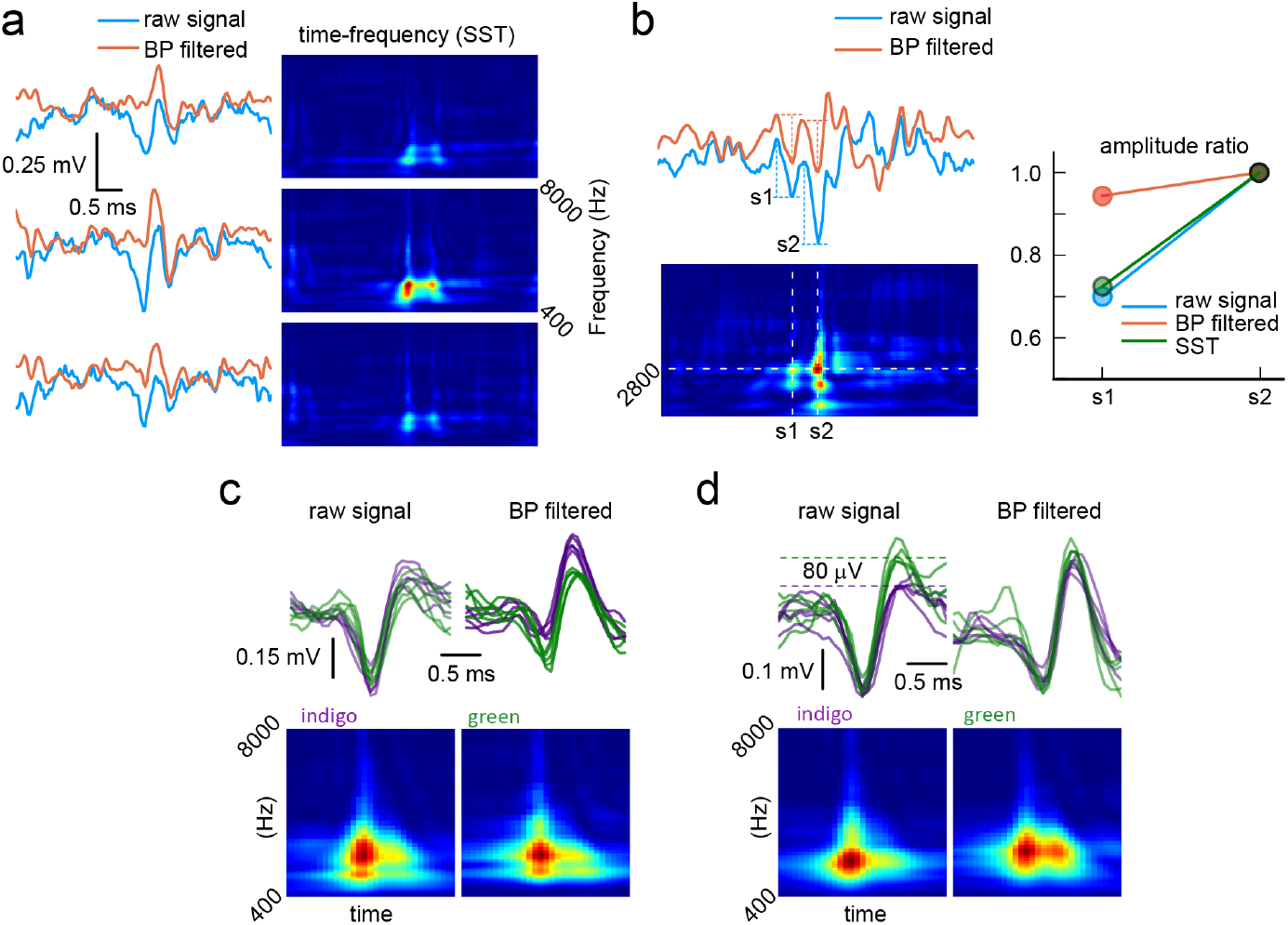
SST spike detection compared to BPF. **a.** Three examples of spike signal disruption caused by band-pass filtering when there is near coincidence of more than one spike signature or other high-frequency event. Note the differences between the blue traces (raw recording) and the orange traces (band-pass filtered), where the latter method will frequently cause full miss of one of the spikes. The three examples represent a signal generated by the same event across three adjacent recording electrodes. As shown in the time-frequency diagrams, the SST method, being based on the raw recording signal, separates the near coincident events in each of the three electrodes. **b.** One example of a spike signature fusion caused by band-pass filtering for near-coincident signals, which the SST method correctly identifies as two different spike signatures. s1 and s2 indicates the times of occurrence of two different spike signatures, reflected in their peak amplitude ratio. The diagram to the right shows that the band-pass filtered signal incorrectly distorts that peak amplitude ratio. **c.** Different occurrences of spikes from the same cluster where the band-pass filtering incorrectly separated the spikes into two different clusters/signatures (spike traces indicated in green and indigo in the upper right). The SST plots at the bottom represent the geometric mean of the SST decompositions across the indigo spikes and the green spikes, respectively. **d.** Fusion of spikes from two different clusters caused by the band-pass filtering. In this case, green and indigo traces represent two spike signatures that are actually different. However, band-pass filtering (upper right) incorrectly results in these two signatures being classifiable as one and the same. The SST method (bottom) correctly separates these as two different signatures.

Fig.3a demonstrates how two spiking events, generated by two different neurons with a small separation in time, appears across 3 neighboring electrodes. In the middle, two spiking waveforms are apparent in the raw time-voltage trace (blue), whereas in the band-pass filtered trace (orange) the two spikes have essentially merged into a single signal, which have the appearance of a valid neural spike. Hence, in this case, BPF resulted in a full miss of one of the spike signals. Furthermore, the spike waveform that did emanate after BPF had a different shape than any of the two ‘parent’ spikes and would thus be at risk of being classified as a third spike that would only ‘occur’ when the spikes of the two neurons coincide in time. In contrast, SST correctly isolated the two events, even when limited to the same frequency range as the BPF, while maintaining the fidelity of the information with respect to the original raw signal. We next investigated BPF’s effects on the spike amplitude information, since amplitude is an important component of extracting features, which are then used for assigning a spike to a cluster. Fig.3b illustrates an example of such an effect with two separate spikes on top of a fast field potential. The BPF distorted the amplitude difference between the two spikes, as quantified to the right, whereas SST instead maintained the relative amplitude information between the spikes. In addition, SST provided information indicative of a fast field potential with an onset in between these two spiking events.

We found that BPF also caused other types of information distortion. Fig.3c shows that for a series of raw spike waveforms taken from the same cluster, BPF generated two distinct waveforms, which a principle component analysis [36] or singular value decomposition [37] based classifier would interpret as two separate neuronal units. However, the mean SST data for the two sets of spikes separated by BPF indicated that these two signals generated the same time-frequency components, meaning they were spikes generated by the same neuron, similarly to what the raw data suggests. Fig.3d illustrates that BPF could also inadvertently fuse two separate spike waveforms into one. The lower part of Fig.3d shows that the geometric means of the SST data for the two different clusters maintained the difference between the two waveforms, maintaining the fidelity with the raw data.

### 2.4 Clustering neuronal spikes with SST

In the next analysis, we investigated if SST could provide more specific representations of the spiking waveforms than BPF, i.e. in analogy with the cocktail party problem described in the Introduction, how well these approaches could perform in the voice fingerprint separation (correctly spotting individuals based on the frequency characteristics of their voices).

For this analysis, we used the raw time-voltage recording data across 8 neighboring electrodes (Fig.4a) for 100 spikes each of 5 pre-identified spike clusters, which are shown as overlaid traces of un-filtered and un-denoised data in Fig.4b. Notably, since all spikes were *a priori* given to the classifier, this part of the analysis ignores the fact that BPF would likely have entirely missed subsets of these spikes, as shown in Fig.3.

**Fig. 4.**
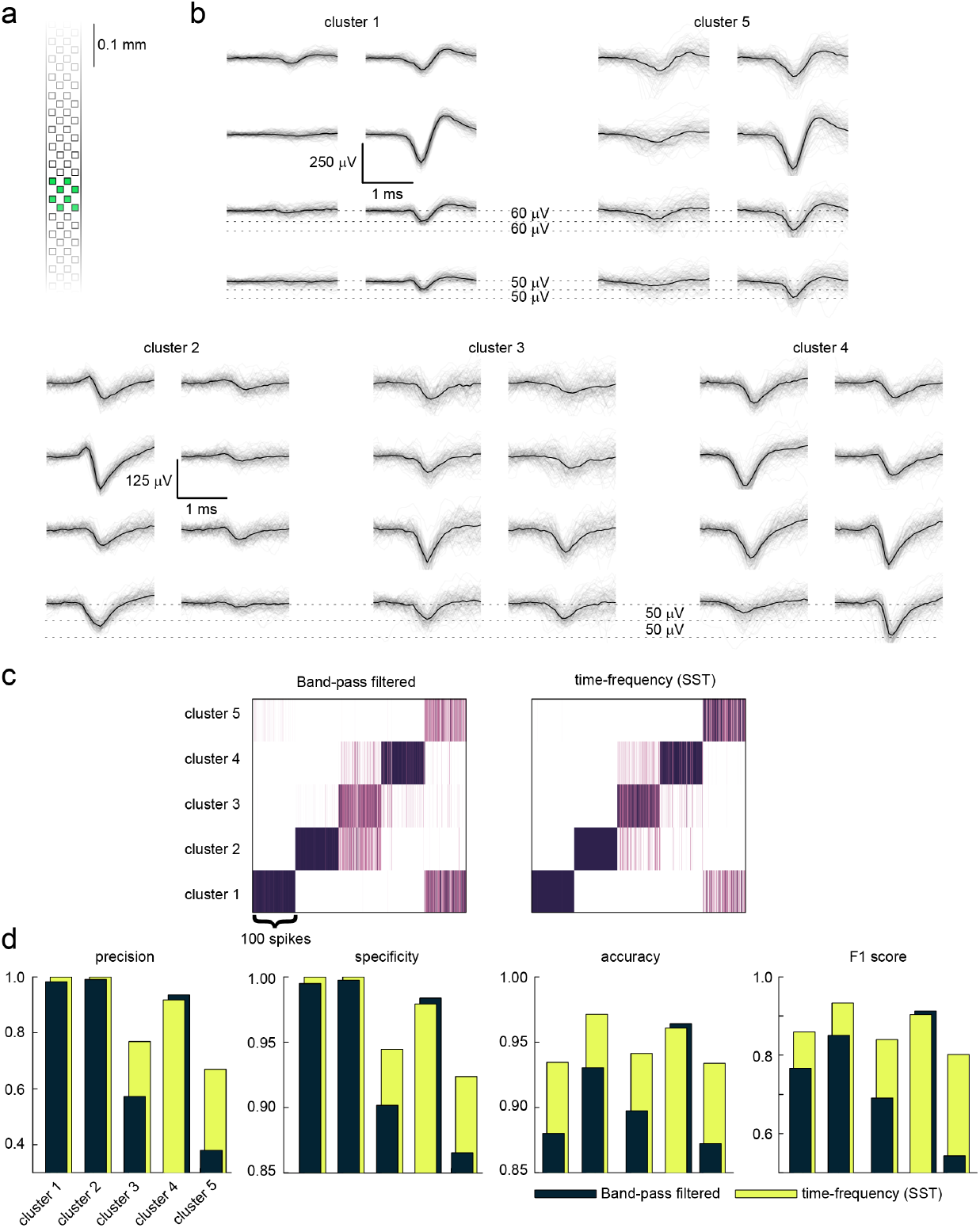
Advantages of SST over BPF in spike clustering. **a.** Sketch indicating the 8 recording electrodes of the NeuroPixel probe used for illustration. **b.** 100 overlaid spike waveforms for 5 identified spike clusters across the 8 electrodes. The median waveform for each electrode is shown in bold. Horizontal lines clarify amplitude differences between subsets of the clusters. **c.** Confusion matrices for the 5 spike clusters generated by spike identification data extracted from the band-pass filtering method and from SST, respectively. **d.** Clustering performance metrics for the two approaches.

For each raw waveform, we used the vectors described by its voltage over time (for BPF) or by its power across specific frequency bands over time (for SST). The classifier then simply compared the mathematical proximity of the resulting vectors for the pool of raw spike waveforms. Notably, whereas each time step (sampling interval) was one dimension per recording electrode for the BPF vector, for SST each time step instead represented information across 50 dimensions, one for each frequency band considered (see Methods), per recording channel. For each spiking waveform we assigned a similarity score based on its most proximal spiking waveforms in the Euclidean space (Fig.4c), which allowed us to quantify the intracluster similarity (precision) and intercluster separability (specificity) of the spike waveform fingerprints (Fig.4d) across all the recorded spikes of the 5 different neurons (Fig.4c,d), for each of the two methods (BPF and SST).

The confusion matrices in Fig.4c show that SST achieved a better clustering than BPF. A particularly difficult case for the classifier was the tendency of confusion between cluster 1 and cluster 5, a confusion which indeed on inspection of the raw data panels in Fig.4b does not appear surprising as these spikes had a similar relative amplitude profile across the 8 electrodes (but note that there were clear pairwise amplitude differences between the signals across 3 of the electrodes, > 10 times the root-mean squared (r.m.s.) of the baseline [3]). In this difficult case, Fig.4c shows that the SST clearly outperformed BPF where the precision went from 38% with BFP to 67% with SST. Hence, almost 1/3rd of all spikes would have went from cluster 5 to cluster 1 with BPF, but were sorted much more accurately into their respective clusters with SST. Similarly in the case of cluster 3, the precision went up from 57% to 76%, corresponding to about 1/5th of all spikes from that cluster ending up in cluster 2 and cluster 4 with BPF, but that were ‘rescued’ with SST. Note that there was again > 10 times the (r.m.s.) pairwise amplitude difference between at least 3 of the electrodes for these 3 clusters (Fig.4b).

### 2.5 Applying SST to audio data

Time-frequency decomposition methods have applicability across a range of domains. Here we provide as an example how the SST method works with sound signals. Fig.5 shows a time-frequency decomposition of a piece of classical music (30 ms of an orchestral performance of Eine Kleine Nachtmusik, between 1.04 and 1.07 s into the piece). The SST output is here compared only with CWT for parameter equivalence, which in turn is analogous in time-frequency representation to STFT (or spectrogram), commonly used in audio decomposition. Notably, despite that the frequency representations were similar, SST achieved a dramatic improvement in the specificity of the temporal representation. Beyond providing advantages for audio decomposition as such, SST could hence for example also provide substantial advantages within the context of auditory neuroscience, with a potential drastic improvement in the precision of temporal correlations between the audio input and the neural responses to it.

**Fig. 5.**
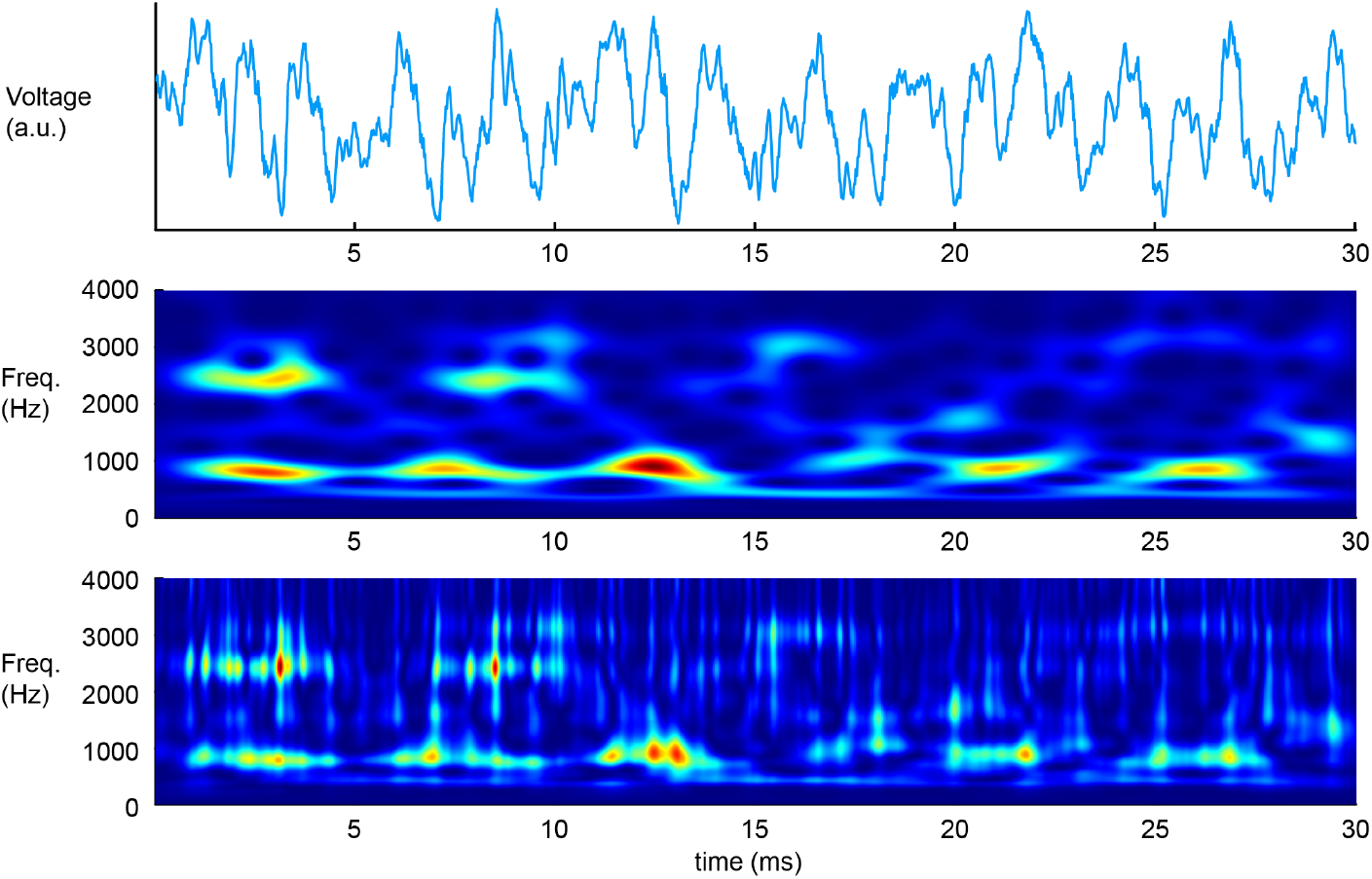
SST achieves more resolvable time-frequency decomposition of music. The voltage signal of a 30 ms sound clip from ‘Eine Kleine Nachtmusik’ and its time-frequency decomposition using CWT (middle) and SST (bottom).

## 3 Discussion

Here we introduced the singular superlet time-frequency decomposition method, and showed that it both increased resolution and reduced computational load compared to the closest comparable super-resolution method, SLT. The singular superlet is analytically combining the effect of multiple discrete observations into a single expression. This resulted in a much reduced computational load for SST (Fig.2b) as well as a much higher resolving power (Fig.1, Fig.2a). The singular superlet is a super-resolution wavelet, where an adaptive adjustment of the number of cycles of the wavelet with respect to frequency achieves the singular superlet transform (SST). We demonstrated the power of the SST in the well-known signal processing problem of unmixing complex recordings containing multiple neuronal spike and field potential signals.

### 3.1 General theoretical considerations

Any time-frequency decomposition is subject to the Heisenberg uncertainty principle, which imposes a fundamental mathematical limitation on the resolving power. This implies that any increase in perceived resolving power will come with a trade-off that may not be perceived in one application, but could be perceived in another. As shown in Fig.2c, the resolving structure of the singular superlet indicates that SST is sacrificing certainty at its tails off the central point of interest (time of occurrence and central frequency, respectively), which can render SST sub-optimal for an application that requires confining long cyclical events in as narrow a window as possible with as high certainty as possible. But the short-burst super-resolution capacity of SST makes it exceptionally useful for analyzing data that is composed of fast and short bursting events, as in the case of neuronal spike recordings in low-impedance extracellular microelectrodes with multiple overlaid signal sources.

### 3.2 Spike clustering

Automated spike sorting of extracellular neural recordings is a problem that is only partially solved, as demonstrated by the number of publications that approach this problem from alternating angles over the years [2, 3, 10, 27–35]. Despite each sorting algorithm approaching the problem from a different angle, filtering have remained a core element in spike sorting. Being able to express the raw signal data with high fidelity at the frequency range of interest, SST can provide a lower distortion filtering (Fig.3) for spike detection and feature extraction, where both the perceived time-frequency resolution improvement 2c and the lower computational load to achieve this super-resolution 2b for the first time (to the best of the authors’ knowledge) allows a time-frequency decomposition method to replace band-pass filtering in the spike sorting pipeline. This is advantageous since a time-frequency decomposition, instead of a directly altering the information content like a BPF, is meant to express the same information content as the raw data with a different vocabulary in order to maintain the fidelity with the original data. Apart from the spike sorting itself, this discrepancy in fidelity can be particularly limiting in determining specific cell types assigned based on their spiking waveforms after band-pass filtering [35, 38].

Notably the difference in fidelity not only demonstrates itself in cases with interferences, but also in spike sorting of *a priori* identified spikes. As demonstrated in figure 4, despite clear amplitude differences across multiple electrodes, BPF can smear the fingerprints of the spikes so much that for 2 out of 5 given neuronal clusters, the miss-rate in precision was as high as 43% and 62%. These misplaced spikes end up in other clusters, which in a sorting algorithm based on BPF could result in neuronal clusters that either contains erroneous firing information, or gets discarded due to high contamination. SST reduces this type of error by nearly half-fold by generating high fidelity representations of the specific spiking fingerprints. This observation implies that when applied to *a priori* extracted spikes, BPF is not as reliable as SST in placing the individual members of the same cluster close to each other (intracluster similarity) despite being as reliable in placing the members of different spike clusters away from each other in Euclidean space (intercluster separability). However, information fed to a clustering algorithm needs to be reliable for both measures.

One possible explanation for this imprecision with intracluster similarity is that BPF is mostly operating by preserving a spike-like event across the electrodes, while the signal fingerprint of the spike waveform, i.e. the detailed information in the frequency domain, has a tendency to be smeared out by the BPF as shown in Fig.3. Then the spike classifier has to rely more on the distribution of the spike amplitudes across the different electrodes to identify a specific neuron, meaning as in the case with cluster 3 and cluster 5 in figure 4, similar amplitude distributions create overlapping information content with other neurons, compromising the precision.

Time-frequency decomposition methods such as SST avoid modifying the raw recording signal information, and instead it uses a different ‘vocabulary’ to express the same information in the frequency range of interest. SST still achieves a ‘BPF-like’ effect by disregarding the low frequency field potentials and the high frequency background noise, while having good enough time resolution for isolating the peaks of the spikes and good enough frequency resolution to preserve the spike features. Looking at the f1 scores (Fig.4d), which is the overall determinant for the reliability of the underlying spike sorting information, SST was only on par with band-pass filtering in one case (cluster 4) and had considerable advantages in every other case, especially when the cases got more challenging (with similar spike amplitude distributions across the electrodes).

The problem of over-merging or over-disregarding neuronal clusters can also manifests itself as significant discrepancies between the number of theoretically available neurons and actually recorded neurons [7, 25] or as a misrepresentation of neural population dynamics. SST can alleviate these problems from the early stages of spike sorting pipeline and thereby both increase the number of neurons detected and the quality of the neuronal clusters created, increasing the reliability of the findings within the field.

### 3.3 Towards an automated spike sorter

Despite our emphasis on inspecting raw biological intracortical data in the context of spike sorting, we’re not proposing this method as a standalone spike sorting algorithm for dense multichannel intracortical recordings. That would require additional components, including but not limited to, automated unsupervised clustering, drift correction, a graphical interface for error correction, etc. that is beyond the scope of this work that aims to present SST as a generic and reliable time-frequency decomposition method that can be utilized as a component for building a solution to the long standing spike sorting problem in neuroscience.

### 3.4 SST is a general signal processing method

Here we presented a new signal processing method, the singular superlet transform (SST). Beyond neuroscience, the generic applicability of SST was presented as an inspection of synthetic test signals and in audio processing. SST drastically improved the resolution in all inspected cases compared to state-of-the-art. We expect SST to find applications within many more domains of signal processing.

## 4 Methods

### 4.1 Synthetic Test Data

Each chirp in the ‘overlaid chirps’ *y* in (fig. 2a) was generated using the following formula:

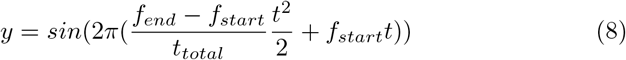

where the time vector *t* is from 0 to *t_total_* = 50 ms sampled at 30 kHz and *f_start_* and *f_end_* correspond to the starting and ending frequencies of the chirps.

### 4.2 Wavelet Normalization

In figure 2a all wavelets (including the singular superlet) used inside a transform are normalized with the following equation:

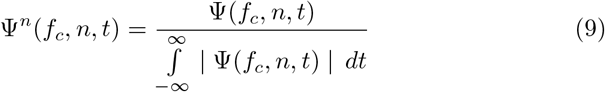

For the rest of the paper, all wavelets (including the singular superlet) used inside a transform are normalized with the following equation:

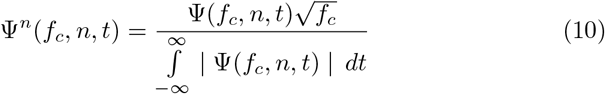

### 4.3 Experimental model and subject details

An adult Sprague-Dawley rat of male sex (weight 320 g) was used in the acute experiment. The animal was maintained in the Lund University animal facilities under 12h light/dark condition. 2–3 rats were habituated in a cage of type 3H, before the experiments, and they were under ad libitum condition to have free access to food and water.

### 4.4 Surgical procedures

In order to make the acute in vivo recording, the adult Sprague Dawley rat was sedated by inhaling air mixed with isoflurane gas (3%, for 2 min). To induce general anesthesia, a mixture of ketamine/xylazine (ketamine: 100 mg/kg and xylazine: 20 mg/kg, accordingly) was injected intraperitoneally. An incision in the inguinal area of the hind-limb was made to insert a catheter in the femoral vein for continuous infusion of Ringer acetate and glucose mixed with anesthetic (ketamine and xylazine in a 20:1 ratio, delivered at a rate of 5 mg/kg/h ketamine). After the initial preparation steps, the somatosensory cortex (SI); was exposed by removing a small part of the skull on the right-hand side (1 × 1 mm), located at (from bregma): Ap: - 1.0 - +0.1, ML: 3.0–5.0. A small incision was made in the skin above the spinotrapezius muscle to attach the grounding electrode.

The level of anesthesia was assessed both by regularly verifying the absence of withdrawal reflexes to noxious pinch of the hind paw and by continuously monitoring the irregular presence of sleep spindles mixed with epochs of more desynchronized activity, a characteristic of sleep [40]. In order to prevent the exposed areas of the cortex from dehydrating, and to decrease the brain tissue movements, a thin layer of agarose (0.5%) was put on the exposed cortical areas, after the insertion of the recording probe. The animal was sacrificed by an overdose of pentobarbital at the end of the experiment.

### 4.5 Extracellular recordings

Electrophysiological recordings were performed using commercially available Neuropixels 1.0 acquisition hardware [3] in conjunction with PXIe-1071 chassis and PXI-6133 I/O from National Instruments and is exported with SpikeGLX software [41]. Sampled at 30k Hz, the amplifier gain used during recording was 500 with the deepest 384 recording sites (‘bank 0’) on the recording probe. Reference and ground was shorted on the ribbon cable and a ground electrode was connected to the integrated printed circuit board on one end, and placed inside the spinotrapezius muscle on the other end.

### 4.6 Estimating spike clustering

For the comparison of the clusterings obtained using SST and band-pass filtering, respectively, we used 0.51 ms vectors (corresponding to 17 sample consecutive points at 30 kHz) of pre-identified spike waveforms. The analysis was based on 100 spikes each of 5 pre-identified spike waveforms. For band pass-filtering, we used a 5th order Butterworth filter between 398 Hz and 7943 Hz. The spikes were aligned based on their peak minimum (and displayed in fig. 4b as aligned), and the time window spanning each spike is centered symmetrically around the peak minimum. For SST time-frequency decomposition of the pre-identified spikes, 50 frequency bands were sampled uniformly between 2.6 to 3.9 raised to the power of 10 (10^2.6^ = 398 Hz, 10^3.9^ = 7943 Hz). From 8 electrode, with 5 clusters of 100 spikes each, this yielded a tensor of [17 × 8 × 500] for band-pass filtered data and [17 × 50 × 8 × 500] for SST data. The proximity of the 500 spikes in Euclidean space were pairwise computed for each method with root mean squared deviation (RMSD), yielding a [500 × 500] matrix per method. Per each of the 500 spikes, a probability estimate was assigned by obtaining the labels for the 51 most proximal spikes (ie: how prevalent each parent cluster label was out of the 51 most proximal spikes) creating a [500 × 5] probability matrix (fig. 4c) where the sum of each [i × 5] column added up to 1.0. In order to go from the per-spike probability scores to cluster metrics (precision, specificity, accuracy and f1 scores, fig. 4d) we averaged the probabilities in each cluster (100 spikes) among themselves, and reduced the values to a [5 × 5] confusion matrix to obtain the true positive, true negative, false positive and false negative rates for each cluster.

## Ethics declarations

Institutional permission The ethical approval for this study was received from the Lund/Malmö local animal ethics committee in advance (permit ID M13193-2017).

## Data Availability

Data will be made available upon request.

## Code Availability

Data were analyzed using custom software. Code will be publicly available upon acceptance.

## Acknowledgments

This research was funded by the EU Horizon 2020 research and innovation program under grant agreement No 861166 (INTUITIVE).

